# Vocal Call Locator Benchmark (VCL) for localizing rodent vocalizations from multi-channel audio

**DOI:** 10.1101/2024.09.20.613758

**Authors:** Ralph E Peterson, Aramis Tanelus, Christopher Ick, Bartul Mimica, Niegil Francis, Violet J Ivan, Aman Choudhri, Annegret L Falkner, Mala Murthy, David M Schneider, Dan H Sanes, Alex H Williams

## Abstract

Understanding the behavioral and neural dynamics of social interactions is a goal of contemporary neuroscience. Many machine learning methods have emerged in recent years to make sense of complex video and neurophysiological data that result from these experiments. Less focus has been placed on understanding how animals process acoustic information, including social vocalizations. A critical step to bridge this gap is determining the senders and receivers of acoustic information in social interactions. While sound source localization (SSL) is a classic problem in signal processing, existing approaches are limited in their ability to localize animal-generated sounds in standard laboratory environments. Advances in deep learning methods for SSL are likely to help address these limitations, however there are currently no publicly available models, datasets, or benchmarks to systematically evaluate SSL algorithms in the domain of bioacoustics. Here, we present the VCL Benchmark: the first large-scale dataset for benchmarking SSL algorithms in rodents. We acquired synchronized video and multi-channel audio recordings of 767,295 sounds with annotated ground truth sources across 9 conditions. The dataset provides benchmarks which evaluate SSL performance on real data, simulated acoustic data, and a mixture of real and simulated data. We intend for this benchmark to facilitate knowledge transfer between the neuroscience and acoustic machine learning communities, which have had limited overlap.

## 1 Introduction

An ongoing renaissance of ethology in the field of neuroscience has shown the importance of conducting experiments in naturalistic contexts, particularly social interactions [38, 1]. Most experiments in social neuroscience have focused on relatively constrained contexts over short timescales, however an emerging paradigm shift is leading laboratories to adopt longitudinal experiments in semi-natural or natural environments [49]. With this shift comes significant data analytic challenges—such as how to track individuals in groups of socially interacting animals—necessitating collaboration between the fields of machine learning and neuroscience [10, 43].

Substantial progress has been made in applying machine vision to multi-animal pose tracking and action recognition [44, 31, 36, 55], however applications of machine audio for acoustic analysis of animal generated social sounds (e.g. vocalizations or footstep sounds) have only recently begun [48, 19]. To study the dynamics of vocal communication and their neural basis, ethologists and neuroscientists have developed a multitude of approaches to attribute vocal calls to individual animals within an interacting social group, however many existing approaches for vocalization attribution necessitate specialized experimental apparatuses and paradigms that hinder the expression of natural social behaviors. For example, invasive surgical procedures, such as affixing custom-built miniature sensors to each animal [16, 47, 60], are often needed to obtain precise measurements of which individual is vocalizing. In addition to being labor intensive and species specific, these surgeries are often not tractable in very small or young animals, may alter an animal’s natural behavioral repertoire, and are not scalable to large social groups. Thus, there is considerable interest in developing non-invasive sound vocal call attribution methods that work off-the-shelf in laboratory settings.

Sound source localization (SSL) is a decades old problem in acoustical signal processing, and several neuroscience groups have adapted classical algorithms from this literature to localize animal sounds [40, 53, 62]. These approaches can work reasonably well in specialized acoustically transparent environments, however they tend to fail in reverberant environments (see Supplement) that are required for next-generation naturalistic experiments.

Data-driven modeling approaches with fewer idealized assumptions may be expected to achieve greater performance [65]. Indeed, in the broader audio machine learning community, deep networks are commonly used to localize sounds [21]. Typically, these approaches have been targeted at much larger environments—e.g. localizing sounds across multi-room home environments [51]. To our knowledge, no benchmark datasets or deep network models have been developed for localizing sounds emitted by small animals (e.g. rodents) interacting in common laboratory environments (e.g. a spatial footprint less than one square meter). To address this, we present benchmark datasets for training and evaluating SSL techniques in reverberant conditions.

## 2 Background and Related Work

### 2.1 Existing Benchmarks

Acoustic engineers are interested in SSL algorithms for a variety of downstream applications. For example, localization can enable audio source separation [34] by disentangling simultaneous sounds emanating from different locations. Other applications include the development of smart home and assisted living technologies [18], teleconferencing [61], and human-robot interactions [32]. To facilitate these aims, several benchmark datasets have been developed in recent years including the L3DAS challenges [22, 23, 20], LOCATA challenge [14], and STARSS23 [51].

Notably, all of these applications and associated benchmarks are (a) focused on a range of sound frequencies that are human audible, and (b) focused on large environments such as offices and household rooms with relatively low reverberation. Our benchmark differs along both of these dimensions, which are important for neuroscience and ethology applications.

Many rodents vocalize and detect sounds in both sonic and ultrasonic ranges. For example, mice, rats, and gerbils collectively have hearing sensitivity that spans ∼50-100,000 Hz with vocalizations spanning ∼100-100,000 Hz [42]. Localizing sounds across a broad spectrum of frequencies introduces interesting complications to the SSL problem. Phase differences across microphones carry less reliable information for higher frequency sounds (see e.g. [27]). Moreover, a microphone’s spatial sensitivity profile will generally be frequency dependent (see microphone specifications for ultrasonic condenser microphone CM16-CMPA from Avisoft Bioacoustics). Therefore, sounds emanating from the same location with the same source volume but distinct frequencies can register with unique level difference profiles across microphones. Thus, different acoustical computations are required to perform SSL for high and low frequency sounds. Indeed, we find that deep networks trained on low frequency sounds in our benchmark fail to generalize when tested on high frequency sounds (see Supplement).

Moreover, many model organisms (rodents, birds, and bats) are experimentally monitored in laboratory environments made of rigid and reverberant materials. The use of these materials is necessary to prevent animals from escaping experimental arenas, which is of particular concern when doing longitudinal semi-natural experiments. For example, in attempts to mitigate reverberance using specialized equipment such as anechoic foam and acoustically transparent mesh, we found that gerbils will climb or chew through material after a short time in the arena. Therefore, use of hard plastic materials, even at the expense of being more reverberant, is required. Thus, the prevalence and character of sound reflections is a unique feature of the VCL benchmark. For variety, we also include benchmark data from an environment with sound absorbent wall material (E3).

### 2.2 Classical work on SSL in engineering and neuroscience

Conventional methods for SSL from acoustic signal processing are summarized in [11]. These methods primarily use differences in arrival times or signal phase across microphones to estimate sources; differences in volume levels are often ignored as a source of information (but see [3]). We use the Mouse Ultrasonic Source Estimation (MUSE) tool [40, 62] as a representative stand-in for these classic approaches in our benchmark experiments. An alternative method based on arrival times was recently proposed by Sterling, Teunisse, and Englitz [54] (see also [41]).

Neural circuit mechanisms of SSL have been extensively studied in model organisms like barn owls, which utilize exquisite SSL capabilities to hunt prey [29]. Neurons in the early auditory system represent both interaural timing and level differences in multiple animal species [9, 4, 7]. Behavioral studies in humans also establish the importance of both interaural timing and level differences [5], and the relative importance of these cues depends on sound frequency and the level of sound reverberation, among other factors [33, 27, 15]. Altogether, the neuroscience and psychophysics literature establishes that animals are adept at localizing sounds in reverberant environments. Moreover, in contrast to many classical SSL algorithms that leverage phase differences across audio waveforms, humans and animals use a complex combination of acoustical cues to localize sounds.

### 2.3 Deep learning approaches to SSL

SSL algorithms account for a variety of event-specific factors including sound frequency, volume, and reverberation. It is challenging to rationally engineer an algorithm to account for all of these factors and the acoustic machine learning community has therefore increasingly turned to deep neural networks (DNNs) to perform SSL. Grumiaux et al. [21] provide a recent and comprehensive review of this literature, including popular architectures, datasets, and simulation methods. Interestingly, researchers have prototyped a variety of preprocessing steps, such as converting raw audio to spectrograms. In our experiments, we apply DNNs with 1D convolutional layers to raw audio waveforms, which are a reasonable standard for benchmarking purposes (see e.g. [59]). Similar to the existing SSL benchmarks listed above, the vast majority of published DNN models have focused on large home or office environments, which differ substantially from our applications of interest.

### 2.4 Acoustic simulations

Across a variety of machine learning tasks, DNNs tend to require large amounts of training data [25]. This is problematic, since it is labor intensive to collect ground truth localization data and curate the result to ensure accurate labels. To overcome this limitation, there is recent interest in leveraging acoustic simulations to supplement DNN training sets. A popular simulation approach is the image source method (ISM) [2]. ISM is computationally inexpensive, relative to alternatives, and preserves much of the spatial information necessary for SSL by accurately modeling the early reflections of sound events. ISM-based approaches to SSL using white noise [8] as well as music, speech and other sound events [30] have shown success when evaluated on data also generated from the ISM.

Recent work has shown that use of room simulations generated using the ISM can also benefit model performance on real-world data [26] and can improve robustness by simulating a wider range of acoustic conditions than is present in an existing training dataset [46], despite perceptual limitations of the ISM. Given these trends in the field, our dataset release includes simulated environments and code for performing ISM simulations.

## 3 The VCL Dataset

The VCL Dataset consists of raw multi-channel audio and image data from 770,547 sound events with ground truth 2D position of the sound event source established by an overhead camera. We recorded synchronized audio (125 or 250 kHz sampling rate) and video (30 Hz or 150 Hz sampling rate) during sound generating events from point sources emanating from either a speaker or real rodents. Sound events were sampled across three environments of varying size, microphone array geometries, and building material (Figure 1A-B, Table 1-2). Ground truth positions were extracted from the video stream using SLEAP [44] or OpenCV, and vocal events from real rodents were segmented from the audio stream using DAS [52]. Timestamps from sound events using speaker playback were either recorded by a National Instruments data acquisition device or pre-computed and used to generate a wav file with known sound event onset times (see Supplement for additional detail).

**Figure 1:**
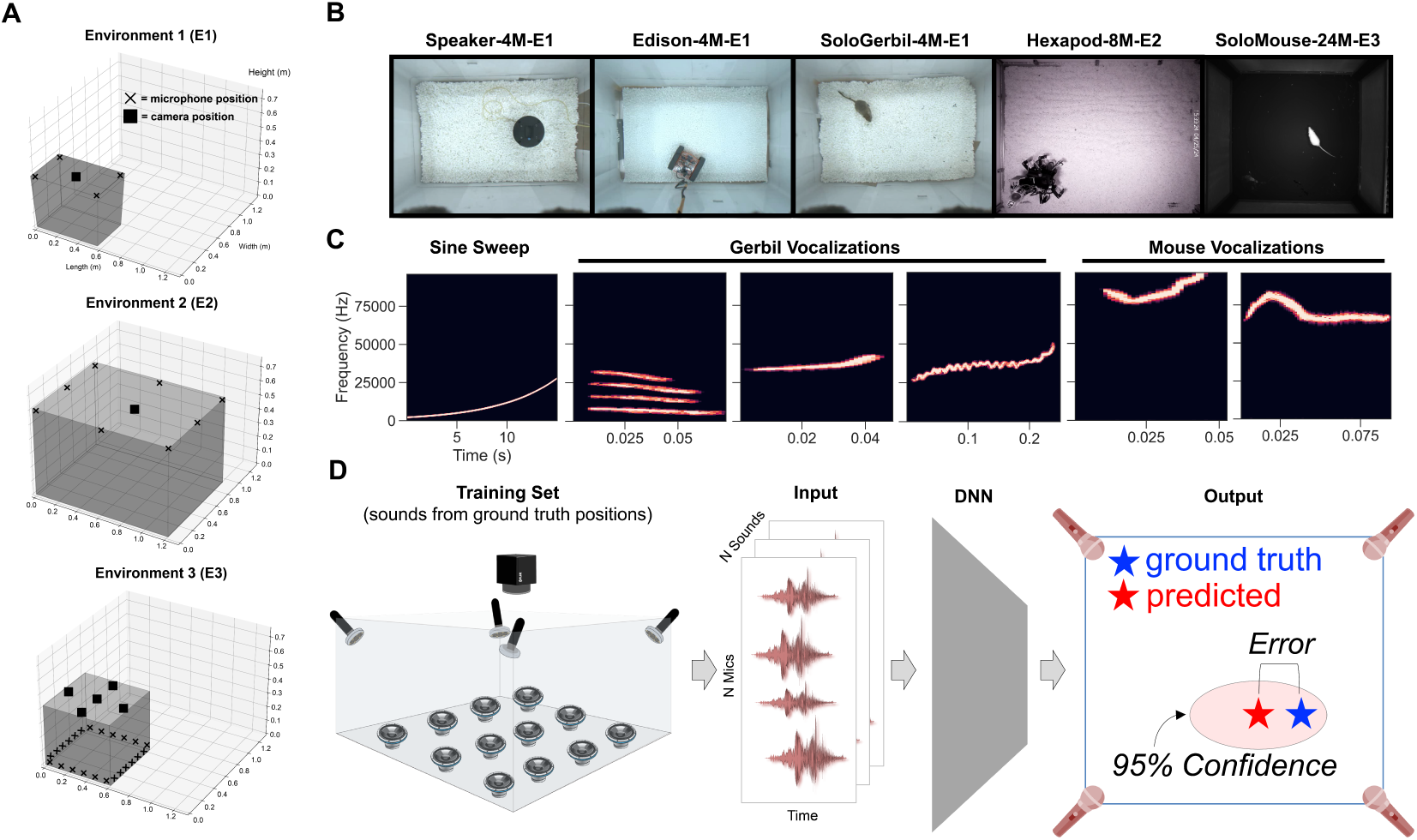
Overview of VCL benchmark. (A) Schematics of three laboratory arenas summarized in Table 2 showing relative size and positions of mics (X’s) and cameras (squares). (B) Top-down views of different environments and training data generation modalities. (C) Examples of stimuli used for playback from Speaker, Edison, Earbud, and Hexapod datasets. (D) Schematic of pipeline depicting inputs (raw audio) and outputs (95% confidence interval).

**Table 1:**
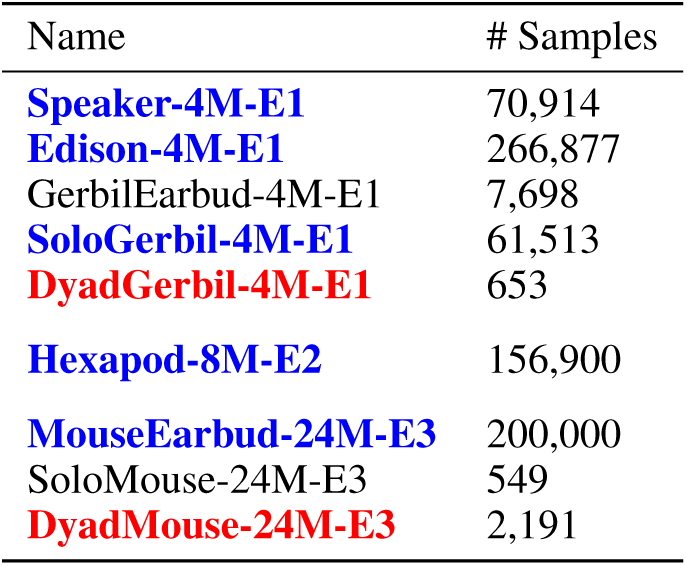
Summary of datasets. Datasets in blue were used as training sets and for test sets when benchmarking SSL. Datasets in red were used as test sets when benchmarking sound attribution.

**Table 2:** Summary of environments. The final two characters in each dataset name (refer to Table 1) specifies the environment in which it was collected.

| Name | # Mics | Dimensions (m) |
| --- | --- | --- |
| E1 | 4 | Top: 0.61595 x 0.41275<br>Bottom: 0.5588 x 0.3556<br>Height: 0.3683 |
| E2 | 8 | Top: 1.2182 x 0.9144<br>Bottom: 1.2182 x 0.9144<br>Height: 0.6096 |
| E3 | 24 | Top: 0.615 x 0.615<br>Bottom: 0.615 x 0.615<br>Height: 0.425 |

For datasets that involved speaker playback, we used primarily rodent vocalizations as stimuli (Figure 1C). In addition, we played sine sweeps in each environment which were used to compute a room impulse response (RIR, see Section 3.5).

### 3.1 Speaker Datasets

The Speaker Dataset (Speaker-4M-E1) was generated by repeatedly presenting five characteristic gerbil vocal calls and a white noise stimulus at three volume levels (18 total stimulus classes) through an overheard Fountek NeoCd1.0 1.5” Ribbon Tweeter speaker. Between every set of presentations, the speaker was manually shifted two centimeters to trace a grid of roughly 400 points along the cage floor. This procedure yielded a dataset of 70,914 presentations spanning the 18 stimulus classes. Gerbil vocalizations can range in frequency from approximately 0.5-60 kHz and different vocalizations correspond to different types of social interactions in nature [57]. In this study, we selected a diverse set of commonly used vocal types vary in frequency range and ethologcial meaning.

### 3.2 Robot Datasets

The generation of the Speaker Dataset was quite labor intensive due to manual movement of the speaker, therefore the procedure was impractical for generating additional training datasets at numerical and spatial scale. To get around this issue, we developed two robotic approaches for autonomous playback of sound events. The Edison and Hexapod Datasets (Edison-4M-E1, Hexapod-8M-E2) were generated by periodically playing vocalizations through miniature speakers affixed to the robots as they performed a pseudo-random walk around the environment. The vocalizations used were sampled from a longitudinal recording of gerbil families [45].

### 3.3 Earbud Datasets

Speaker and robotic playback of vocalizations may not accurately represent the spatial usage and direction of vocalizations in real animals. To address this, we acquired two “Earbud” datasets (GerbilEarbud-4M-E1, MouseEarbud-24M-E3), in which gerbils or mice freely explored their environment with an earbud surgically affixed to their skull. We then played species typical vocalizations out of the earbud while animals exhibited a range of natural behaviors.

### 3.4 Solo/Dyad Gerbil & Mouse Datasets

Although isolated animals usually do not vocalize, we found that adolescent gerbils produce antiphonal responses to conspecific vocalizations played through a speaker. We leveraged this behavior to generate a large scale dataset, SoloGerbil-4M-E1, containing real gerbil-generated vocalizations in isolation. In addition, we elicited solo vocalizations in male mice (SoloMouse-24M-E3) by allowing female mice in estrus to explore the environment prior to male exploration.

Our ultimate goal is to use sound source estimates to attribute vocalizations to individuals in a group of socially interacting animals. To this end, we acquired vocalizations from pairs of interacting gerbils and mice (DyadGerbil-4M-E1, DyadMouse-24M-E3). Although we are unable to determine the ground truth position of vocalizations recorded from these interactions, we do know the locations of both potential sources and can therefore ascertain whether our model generates predictions with zero, one, or two animals within its confidence interval (See Task 2 below).

### 3.5 Synthetic Datasets

Since DNNs often require large training datasets and generation of datasets in the domain of SSL is laborious, we explored the use of acoustic simulations for supplementing real training data (Figure 2). We generated *in silico* models of our three environments accounting for physical measurements of the geometry, microphone placement, microphone directivity, and estimates of the material absorption coefficients (calculated via the inverse Sabine formula on room impulse response measurements with a sine sweep excitation). Code to reproduce these simulations and adapt them to new environments is included in our code package accompanying the VCL benchmark. In preliminary experiments, we found that training DNNs on mixtures of real and simulated data can benefit performance (Figure 2D), but we do not include simulated data in our benchmark experiments described below.

**Figure 2:**
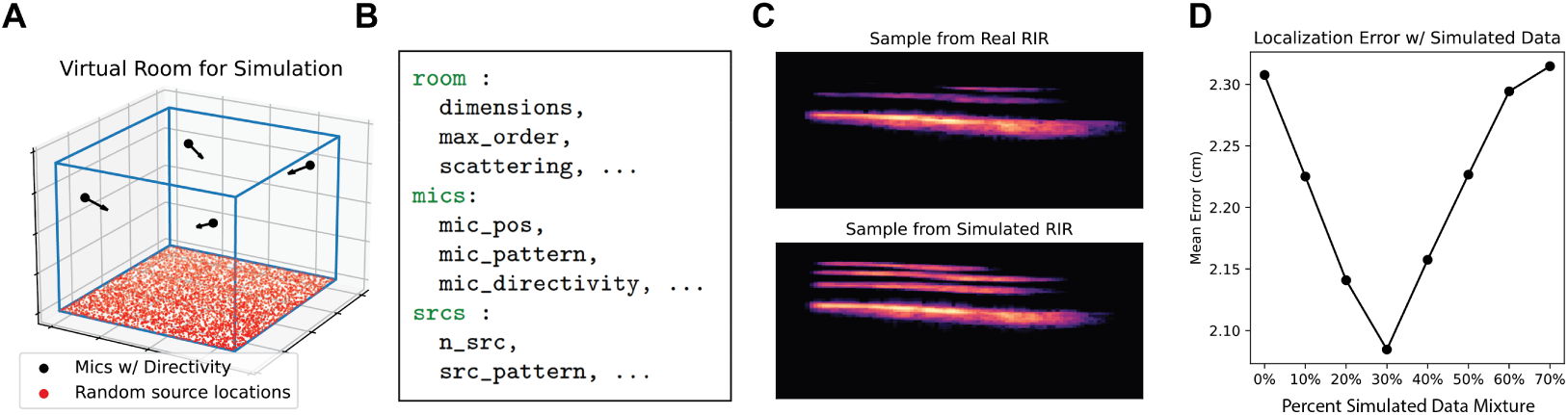
(A) Visualization of virtual room used for sythetic RIR generation via ISM (B) Sample of a room configuration YAML used to specify room geometry for simulations (C) Spectrograms comparing vocalizations convolved with recorded RIRs and simulated RIRs (D) Localization error as a function of added simulated data to the training corpus.

## 4 Benchmarks on VCL

We established a benchmark on the VCL Dataset using two distinct tasks.

- **Task 1 - Sound Source Localization:** Compare the performance of classical sound source localization algorithms with deep neural networks.
- **Task 2 - Vocalization Attribution:** Assign vocalizations to individuals in a dyad.

We evaluated performance on Task 1 using datasets with a single sound source (marked in **blue** in Table 1). We calculated the centimeter error between ground truth and predicted positions. Our aim is to achieve errors less than or equal to ^_^1 cm, as this is the approximate resolution required to attribute sound events to individual animals.^3^ We also sought to benchmark the accuracy of model-derived confidence intervals. That is, for each prediction the model should produce a 2D set that contains the sound source with specified confidence (e.g. a 95% confidence set fail to contain the true sound source on only 5% of test set examples). Following procedures from Guo et al. [24], we plot reliability diagrams and report the expected calibration error (ECE) and maximal calibration error (MCE).

We evaluated performance on Task 2 using datasets with two potential sound sources (marked in **red** in Table 1). For Task 2, we report the number of animals inside the 95% confidence set of model predictions. For each sound event, the model can predict zero, one, or two animals within its confidence set. We report the frequency of each of these outcomes and interpret them as follows. First, if only one animal is within the confidence set, the model attributes the vocalization to that animal. We cannot for verify whether this attribution is correct because (unlike the datasets used in Task 1) we do not have ground truth measurements of the sound source. Second, if two animals are within the confidence set, then the model is unable to reliably attribute the sound to an individual. This outcome is neither correct nor incorrect. Finally, if zero animals are within the confidence set, then the model has falsely attributed the sound to a region. This outcome is clearly incorrect and should ideally happen less than 5% of the time when using a 95% confidence set.

### 4.1 Convolutional Deep Neural Network

The network consists of 1D convolutional blocks connected in series. The network takes in raw multi-channel audio waveforms and outputs the mean and covariance of a 2D Gaussian distribution over the environment. Intuitively, the mean represents the network’s best point estimate of the sound source and the scale and shape of the covariance matrix corresponds to an estimate of uncertainty. The network is trained with respect to labeled 2D sound source positions to minimize a negative log likelihood criterion—this is a proper scoring rule [17] which encourages the model to accurately portray its confidence in the predicted covariance. That is, the 95% upper level set of the Gaussian density should ideally act as a 95% confidence set. However, in line with previous reports, we sometimes observe that DNN confidence intervals are overconfident. In these cases, we use a temperature scaling procedure to calibrate the confidence intervals [24]. Further details on data preprocessing, model architecture, training procedure are provided in the Supplement.

### 4.2 MUSE Baseline Model

We compare the DNNs to a delay-and-sum beamforming approach used by neuroscientists called MUSE [40, 62]. MUSE works by computing cross-correlation signal between all pairs of microphone signals across hypothesized sound source locations, using the distance between microphones and the speed of sound to compute arrival time delays. The location that maximizes the summed response power over all microphones is then selected as a point estimate. We generate 95% confidence sets using a jackknife resampling technique proposed in Warren, Sangiamo, and Neunuebel [62].

### 4.3 Task 1 Results

Deep neural networks consistently produced estimates closer to the ground truth source than MUSE (Figure 3 A-E, Table 3). DNN performance was particularly strong on the Edison-4M-E1 and Speaker-4M-E1 datasets, achieving *<*1 cm error on 80.6% and 66.0% on the respective test sets. As mentioned above, this level of resolution should enable attribution of most vocalizations in realistic social encounters in rodents [54]. DNNs also outperformed MUSE on the remaining three datasets; however, they achieved sub-centimeter errors on less than 10% of the test set in all cases.

**Figure 3:**
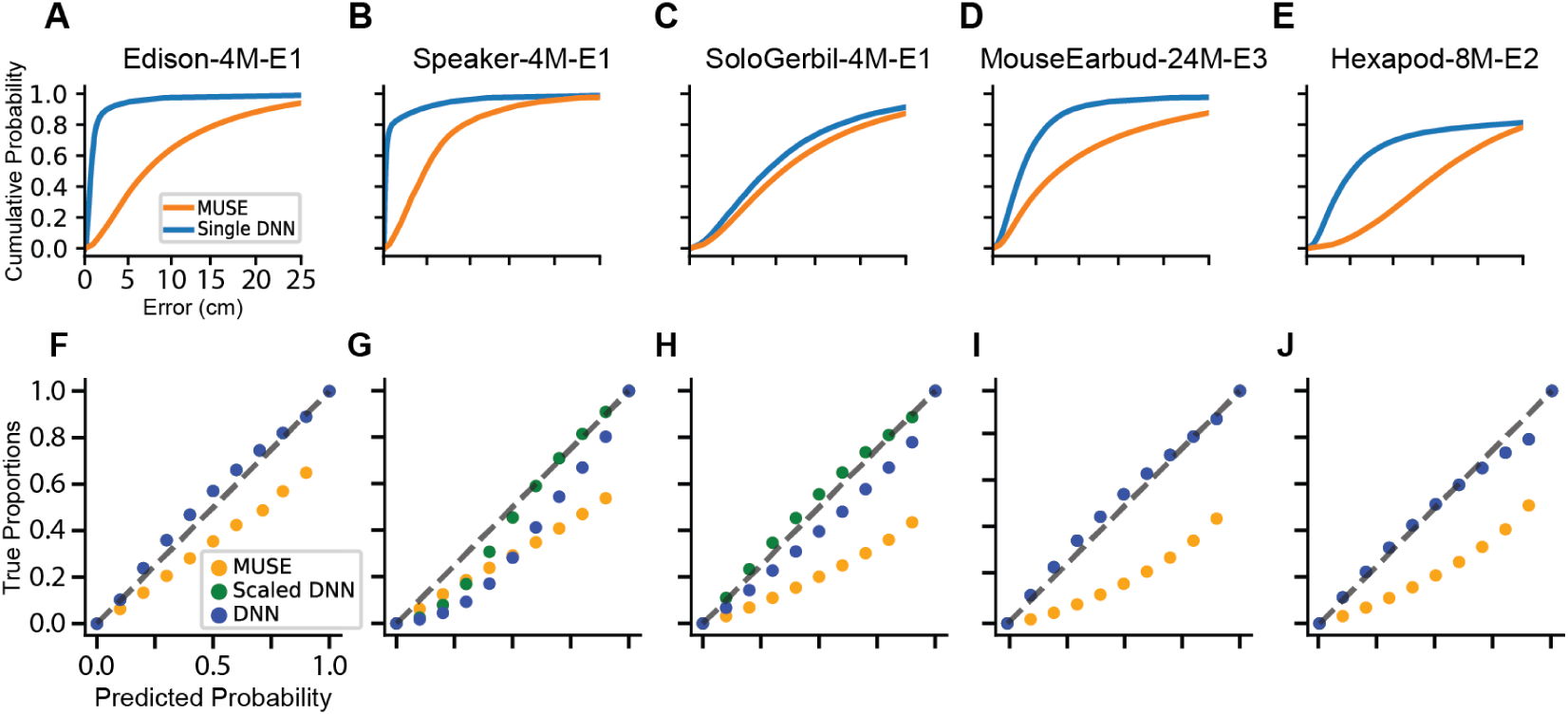
Benchmark performance. (A-E) Cumulative error distributions for MUSE and neural networks. (F-J) Reliability diagrams for MUSE (orange) and neural networks with (green) and without (blue) temperature scaling on heldout data from each dataset.

**Table 3:**
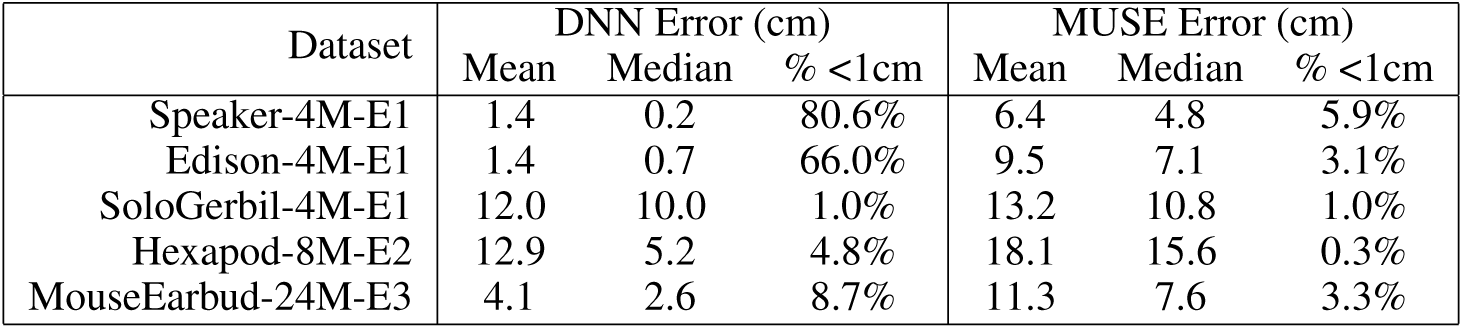
Summary of sound source localization errors for Task 1.

| Dataset | DNN Error (cm) |  |  | MUSE Error (cm) |  |  |
| --- | --- | --- | --- | --- | --- | --- |
|  | Mean | Median | % <1cm | Mean | Median | % <1cm |
| Speaker-4M-E1 | 1.4 | 0.2 | 80.6% | 6.4 | 4.8 | 5.9% |
| Edison-4M-E1 | 1.4 | 0.7 | 66.0% | 9.5 | 7.1 | 3.1% |
| SoloGerbil-4M-E1 | 12.0 | 10.0 | 1.0% | 13.2 | 10.8 | 1.0% |
| Hexapod-8M-E2 | 12.9 | 5.2 | 4.8% | 18.1 | 15.6 | 0.3% |
| MouseEarbud-24M-E3 | 4.1 | 2.6 | 8.7% | 11.3 | 7.6 | 3.3% |

Moreover, we found that DNNs provide more accurate estimates of uncertainty relative to MUSE, as calculated by ECE and MCE (Table 4). This performance difference is visible in reliability diagrams, which show that MUSE predictions are over-confident (Figure 3F-J).

**Table 4:**
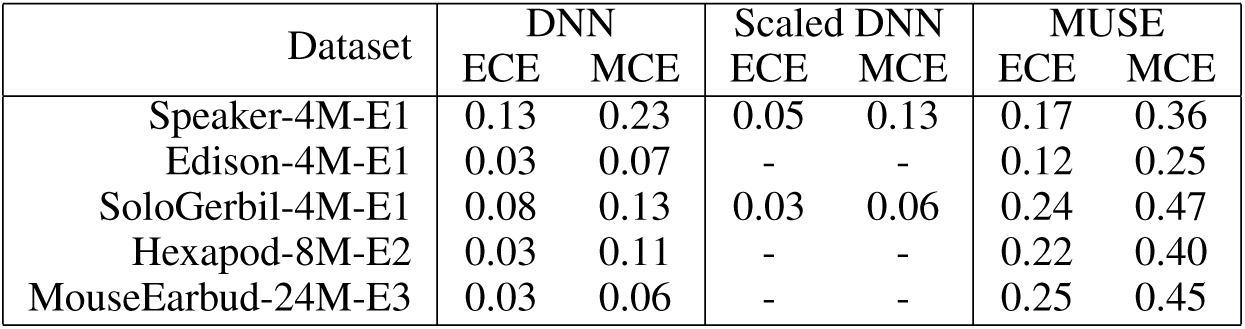
Expected Calibration Error (ECE) and Maximum Calibration Error (MCE) for Task 1.

| Dataset | DNN |  | Scaled DNN |  | MUSE |  |
| --- | --- | --- | --- | --- | --- | --- |
|  | ECE | MCE | ECE | MCE | ECE | MCE |
| Speaker-4M-E1 | 0.13 | 0.23 | 0.05 | 0.13 | 0.17 | 0.36 |
| Edison-4M-E1 | 0.03 | 0.07 | - | - | 0.12 | 0.25 |
| SoloGerbil-4M-E1 | 0.08 | 0.13 | 0.03 | 0.06 | 0.24 | 0.47 |
| Hexapod-8M-E2 | 0.03 | 0.11 | - | - | 0.22 | 0.40 |
| MouseEarbud-24M-E3 | 0.03 | 0.06 | - | - | 0.25 | 0.45 |

### 4.4 Task 2 Results

To test the ability of our DNNs to assign vocalizations to individuals in dyadic interactions, we used DNNs trained on single-agent datasets, MouseEarbud-24M-E3 and SoloGerbil-4M-E1 respectively, to compute confidence bounds on vocalizations from the dyadic datasets MouseDyad-24M-E3 and GerbilDyad-4M-E1. As described above, we used temperature rescaling to ensure DNN confidence sets were well-calibrated. While we were capable of assigning between 19-29% of these calls to a single animal, over half of the vocalizations in each interaction yielded a confidence bound containing both animals (Table 5). Methods to resolve these shortcomings remain a focus of future work.

**Table 5:** Vocalization attribution results. Number of animals captured within the 95% confidence set.

| # Animals Captured | Gerbil Dyad |  |  | Mouse Dyad |  |  |
| --- | --- | --- | --- | --- | --- | --- |
|  | 0 | 1 | 2 | 0 | 1 | 2 |
| Percentage | 6.1% | 28.6% | 65.2% | 8.9% | 19.4% | 71.7% |

## 5 Limitations

Neuroscientists are interested in localizing sounds across a broad range of settings. We aimed to cover multiple rodent species (gerbils and mice), environment sizes, and microphone array geometries in this initial release. We also leveraged robots and head-mounted earbud speakers to collect sounds with known ground truth. However, this benchmark does not yet cover all use cases in neuroscience. Other commonly used model species—e.g., marmosets[13], bats[58], and various bird species[6]—are of great interest and are not covered by the current benchmark. Our experiments show that deep neural networks trained to localize sounds can fail to generalize across vocal call types (see Supplement). It would therefore be valuable to expand this benchmark to include a wider variety of animal species, call types, and increase the number of training samples. To this end, we include additional datasets which were not used in Task 1 due to their relatively small size (GerbilEarbud-4M-E1, SoloMouse-24M-E3), which will aid future experiments assessing generalization performance across datasets (e.g. train on Speaker-4M-E1, predict on GerbilEarbud-4M-E1).

Our current benchmark only provides images from a single camera view, which can be used to localize sounds in 2D. While this agrees with current practices within the field [40, 53, 37] and is in line with the equipment readily available to most labs, it is insufficient to infer 3D body pose information. One could imagination that knowing the 3D position and 3D heading direction of a vocalizing rodent could provide a more rich and effective supervision signal to train a deep network. A number of 3D pose tracking tools for animal models have been developed in very recent years [64, 39, 28, 35, 12]. These tools could be leveraged if future benchmarks collect multiple camera views. Ultimately, it would be useful to compare performance across 3D and 2D benchmarks, to ascertain whether the sound source localization problem is indeed easier in one or the other setting.

## 6 Discussion

SSL is a well-known and challenging problem. We collected a variety of datasets and developed benchmarks to assess these challenges in the context of neuroethological experiments in vocalizing rodents. This involves localizing sounds in reverberant environments across a very broad frequency range (including ultrasonic events), distinguishing our work from more standard SSL benchmarks and algorithms. Our experiments reveal that DNNs are a promising approach. In controlled settings (Edison-4M-E1 and Speaker-4M-E1 datasets), DNNs achieved sub-centimeter resolution. In larger environments (Hexapod-8M-E2) and in datasets with uncontrolled 3D variation in sound emissions (SoloGerbil-4M-E1 and MouseEarbud-24M-E3), DNN performance was less impressive, but still outperformed a well-established benchmark algorithm (MUSE), that is currently utilized.

In addition to continuing to experiment with advances in machine vision/audio, we are also interested in exploring performance improvements due to hardware optimization. Parameters such as number of microphones, their positions/directivity, and environment reverberance can all affect SSL performance. Future experiments will leverage acoustic simulations to explore this parameter space. Initial results suggest that varying the amount of reverberation in an environment drastically affects SSL performance and that this effect is more pronounced in MUSE than DNNs (see Supplement).

The ultimate goal of most neuroscientists is to attribute vocal calls to individuals amongst an interacting social group. Accurate SSL would enable this, but it is also possible to reframe this problem as a direct prediction task. Specifically, given a video and audio recording of *K* interacting animals with ground truth labels for the source of each sound event, DNNs could be trained to perform *K*-way classification to identify the source. Future work should investigate this promising alternative approach, as it would enable DNNs to jointly leverage information from audio and video data as network inputs. On the other hand, we note several challenges that must be overcome. First, establishing ground truth in multi-animal recordings is non-trivial, though feasible in certain experiments [16, 47, 60]. Second, DNNs trained to process raw video can have trouble generalizing across recording sessions due to subtle changes in lighting or animal appearance [63, 50]. Finally, we note that at least *K* = 2 animals are required to make the problem nontrivial (when *K* = 1 the DNN could ignore the audio input to predict the source). It will be important to establish a flexible DNN architecture that can make accurate predictions even when the animal group size, *K*, is altered (see e.g. [66]). It is already possible to use the VCL datasets to explore these possibilities. For example, one could use audio and video data taken from the same or different sound events to train a DNN with a multimodal contrastive learning objective (see e.g. [56], for a related concept).

In summary, there are many promising, but under-investigated, machine learning methodologies for annotating vocal communication in rodents. The VCL benchmark is our attempt to spark a broader community effort to investigate the potential of these computational approaches. Indeed, collecting and curating these datasets is labor-intensive and in our case involved collaboration across multiple neuroscience labs. To our knowledge, very little (if any) comparable data containing raw audio and video from many thousands of rodent vocal calls currently exists in the public domain. Thus, we expect the VCL benchmark will enable new avenues of research within computational neuroscience.

## Acknowledgements and Ethics Statement

We do not foresee any negative societal impacts arising from this work. We thank Megan Kirchgessner (NYU), Robert Froemke (NYU), and Marcelo Magnasco (Rockefeller) for discussions and suggestions regarding SSL applications in neuroscience. This work was supported by the National Institutes of Health R34-DA059513 (AHW, DHS, DMS), National Institutes of Health R01-DC020279 (DHS), National Institutes of Health 1R01-DC018802 (DMS, REP), National Institutes of Health Training Program in Computational Neuroscience T90DA059110 (REP), New York Stem Cell Foundation (DMS), CV Starr Fellowship (BM), EMBO Postdoctoral Fellowship (BM), National Science Foundation Award 1922658 (CI).

## Checklist

1. For all authors…

a. Do the main claims made in the abstract and introduction accurately reflect the paper’s contributions and scope? [Yes]
b. Did you describe the limitations of your work? [Yes] See Section 5.
c. Did you discuss any potential negative societal impacts of your work? [Yes] See Acknowledgements and Ethics Statement.
d. Have you read the ethics review guidelines and ensured that your paper conforms to them? [Yes]
2. If you are including theoretical results…

a. Did you state the full set of assumptions of all theoretical results? [N/A]
b. Did you include complete proofs of all theoretical results? [N/A]
3. If you ran experiments (e.g. for benchmarks)…

a. Did you include the code, data, and instructions needed to reproduce the main experimental results (either in the supplemental material or as a URL)? [Yes] See data website, “vocalator” GitHub repo for DNNs, and supplement.
b. Did you specify all the training details (e.g., data splits, hyperparameters, how they were chosen)? [Yes] See supplement.
c. Did you report error bars (e.g., with respect to the random seed after running experiments multiple times)? [No] Due to time constraints and anecdotal evidence that models from different random seeds produce similar results, we did not include error bars in this draft. We are happy to include them upon revision.
d. Did you include the total amount of compute and the type of resources used (e.g., type of GPUs, internal cluster, or cloud provider)? [Yes] See supplement.
4. If you are using existing assets (e.g., code, data, models) or curating/releasing new assets…

a. If your work uses existing assets, did you cite the creators? [N/A]
b. Did you mention the license of the assets? [N/A]
c. Did you include any new assets either in the supplemental material or as a URL? [Yes]
d. Did you discuss whether and how consent was obtained from people whose data you’re using/curating? [N/A]
e. Did you discuss whether the data you are using/curating contains personally identifiable information or offensive content? [N/A]
5. If you used crowdsourcing or conducted research with human subjects…

a. Did you include the full text of instructions given to participants and screenshots, if applicable? [N/A]
b. Did you describe any potential participant risks, with links to Institutional Review Board (IRB) approvals, if applicable? [N/A]
c. Did you include the estimated hourly wage paid to participants and the total amount spent on participant compensation? [N/A]

## 1 Supplemental Information and Results

### 1.1 Generalization of DNN to heldout predictions

**Figure 1:**
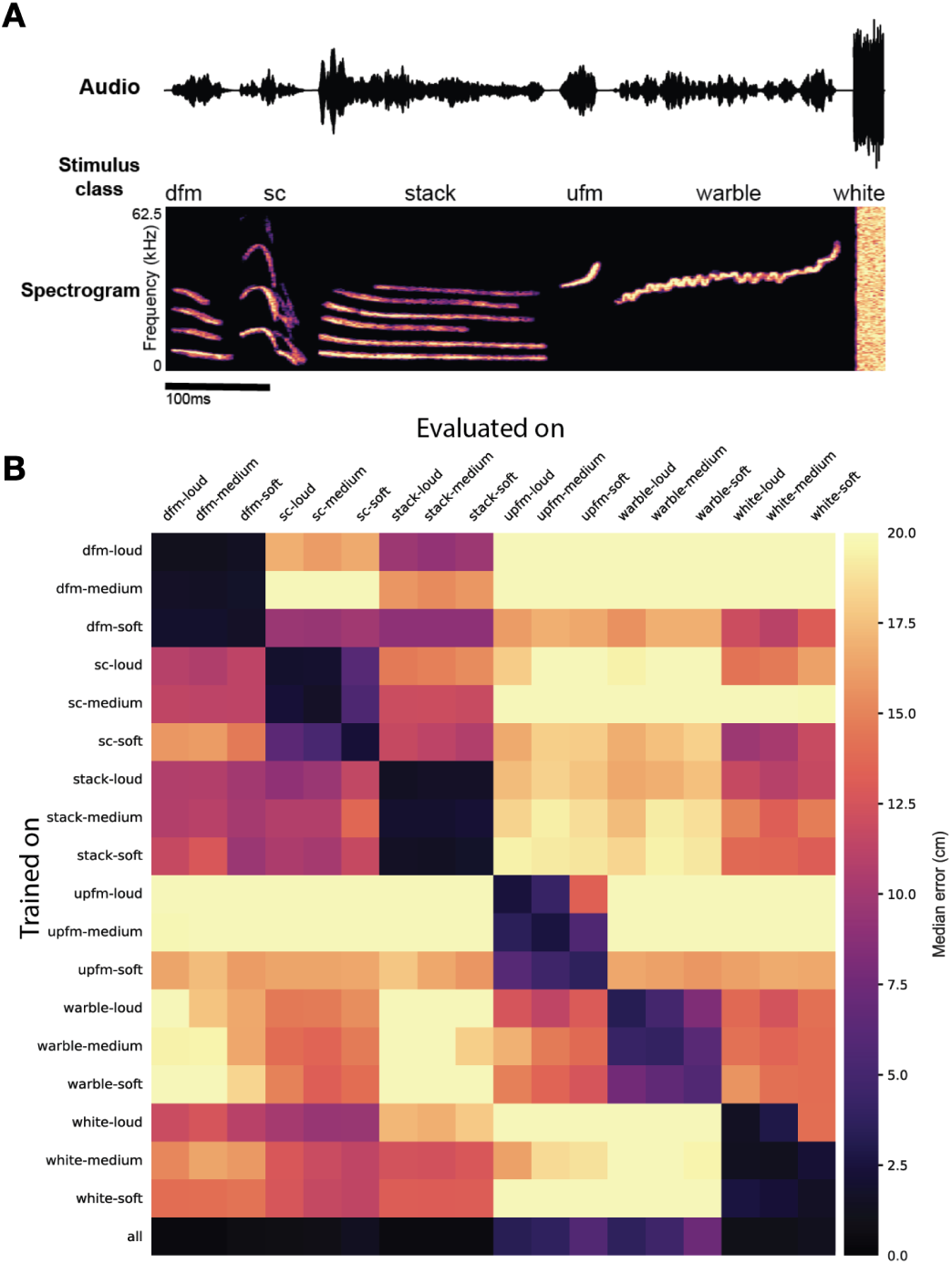
Generalizability across stimulus types. A.) Stimuli used for speaker data set (dfm = down frequency modulated, sc = soft chirp, stack = harmonic stack, ufm = up frequency modulated) B.) Performance of models trained on single stimuli from Speaker-4M-E1 dataset and evaluated on all other stimulus types.

We aim to create a tool that can be easily adapted by other labs which may have different recording environments. Additionally, we wish to utilize the tool for long-term recordings in which the types of vocalizations encountered may change over time as the animals enter new stages of life. As such, we have significant interest in the model’s ability to generalize to unfamiliar vocal calls.

To explore this, we tested the ability of deep networks to generalize to new vocal calls with different acoustic features. We partitioned the Speaker-4M-E1 Dataset according to stimulus type (Supplementary Figure 1A), trained a deep neural network on each subset, and measured its performance on every stimulus type individually (Supplementary Figure 1B). We found that while many models could generalize to new stimuli with performance exceeding chance, their ability to do so is greatly overshadowed by their performance on their own subsets. Models trained on a single stimulus type generalized well to the same stimulus at different volumes. (Supplementary Figure 1B, 3x3 block structure). This suggests that the networks are adapted to the statistics of the training set, and that training on a range of vocalizations with diverse spectral features will be necessary to achieve good performance across experimental cohorts, each of which may utilize slightly different vocal calls.

### 1.2 DNN architecture and training procedure

**Figure 2:**
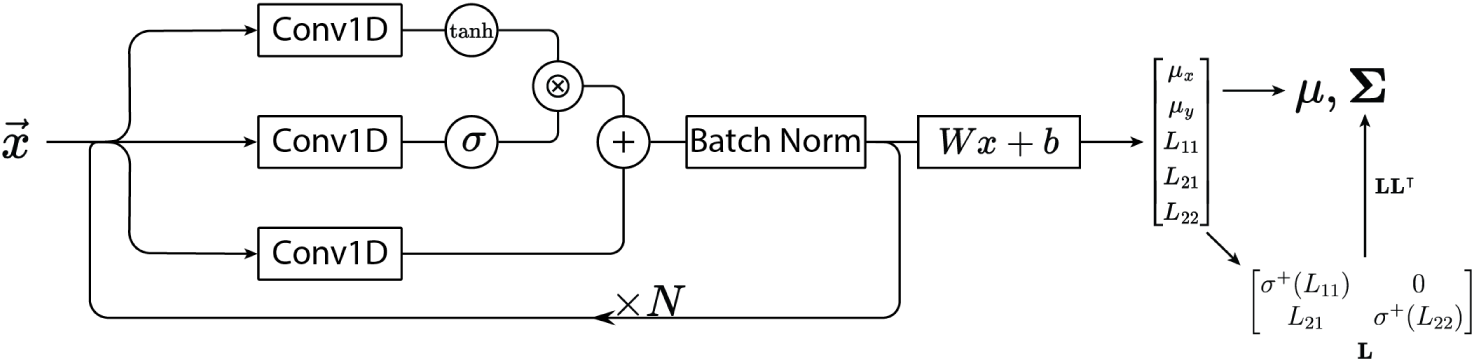
Network architecture.

**Table 1:** Model architecture hyperparameters.

| Layer | Channels | Downsample |
| --- | --- | --- |
| 1 | 32 | No |
| 2 | 32 | Yes |
| 3 | 64 | No |
| 4 | 64 | Yes |
| 5 | 128 | No |
| 6 | 128 | Yes |
| 7 | 256 | No |
| 8 | 256 | Yes |
| 9 | 512 | No |
| 10 | 512 | Yes |

Mirroring gated linear units [1] and WaveNet [9], we apply tanh and sigmoid nonlinearities to the output of convolutions and multiply them element-wise (Supplementary Figure 2). We add this product to the result of a third convolution and apply batch normalization to the sum. On layers with temporal downsampling, we perform average pooling with a stride and kernel size of 2 prior to normalization. On our datasets with four microphones, we incorporate pairwise cross-correlations of the microphone signals by concatenating the central elements of each cross-correlogram into a vector, passing it through a shallow MLP, and concatenating the result to the output of the final convolutional block. The model outputs the mean and covariance of a 2D Gaussian distribution with covariance specified by a Cholesky factor matrix. To parametrize the 2D gaussian, we first average the output of the final convolutional block over its time dimension and linearly project it to five dimensions. Two of these determine the distribution’s mean and the other three parametrize the Cholesky decomposition of the distribution’s covariance matrix. In order to ensure the Cholesky factor has positive diagonals, we apply the softplus nonlinearity to the diagonal elements. During training, we evaluate the log likelihood of the ground truth positions with respect to the 2D Gaussians output by the network. We minimize the negative log likelihood using stochastic gradient descent with momentum. Throughout 50 epochs, we anneal the learning rate to 0 using a cosine schedule. We do not use weight decay. Our model consists of 10 convolutional blocks (Supplementary Table 1). All use a kernel size of 33, dilation of 1, and stride of 1

For data preprocessing, we normalize the audio by ensuring a zero mean and unit variance across all elements, rather than scaling each channel individually. This approach ensures amplitude differences between channels are preserved after normalization. Throughout training, we apply various augmentations to the audio to enhance sample efficiency and performance on the validation set. As vocalization lengths vary substantially, we randomly crop them to a standardized length of 8192 samples (65.5ms at 125kHz) to facilitate batched computations. Additional augmentations include temporal masking, the introduction of white noise, and phase inversion. With the exception of cropping, which is applied universally to all samples, each augmentation has a 50% chance of being applied to a given vocalization.

### 1.3 Impact of reverberation in simulated datasets

**Figure 3:**
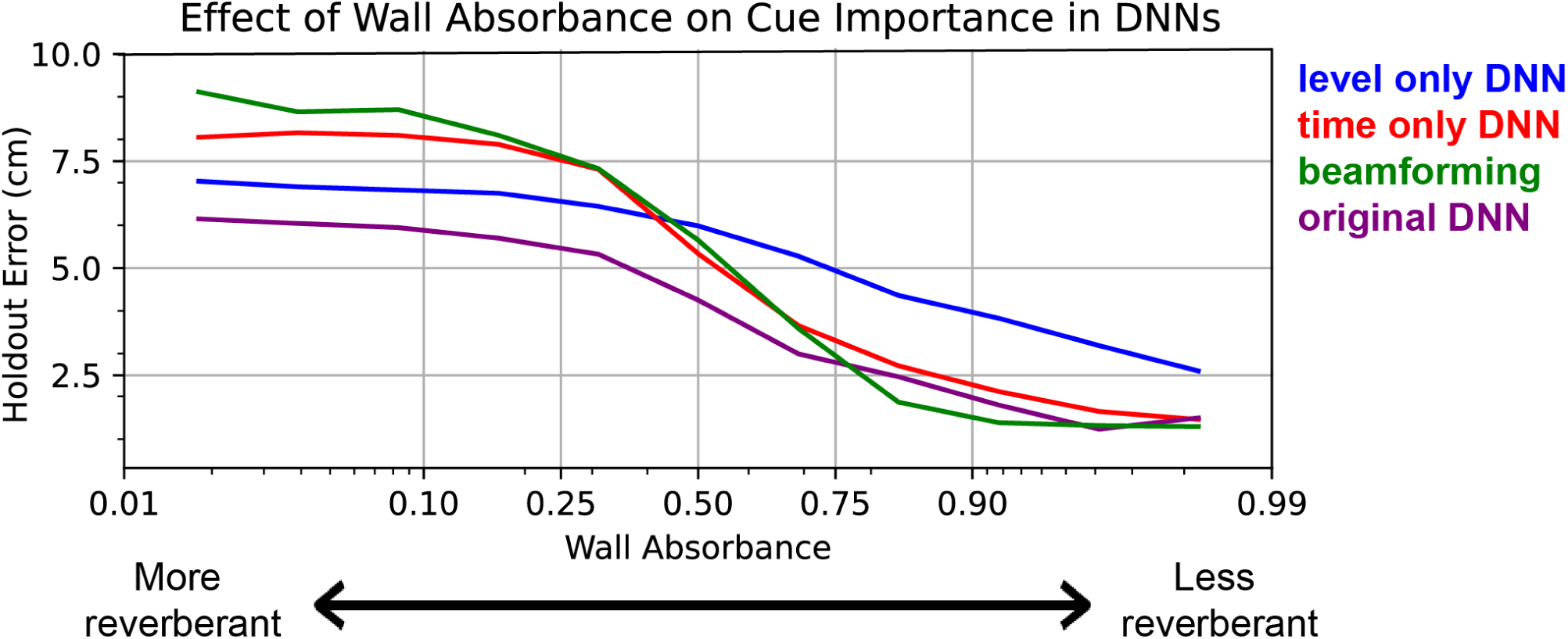
SSL performance on simulated data from the E1 environment with varying environmental reverberance.

We explored whether SSL performance systematically varied as a function of reverberance using acoustic simulations. First, we simulated an E1 environment, then simulated microphone signals from 50,000 gerbil vocalizations randomly sampled from [7]. Next, we compared DNN vs. MUSE (beamforming) performance and showed that DNNs (purple) outperform MUSE (green) in reverberant conditions and achieve equal performance in non-reverberant conditions. Furthermore, we explored which cues (temporal or level, i.e. akin to ITD and ILD cues used by animals) DNNs relied on for SSL. We created augmented training sets that either scrambled level differences between microphone channels (thereby only maintaining reliable time differences, red) or scrambled time differences (thereby only maintaining reliable level differences, blue). We find that time-only DNN performance matches MUSE, which is consistent with the fact that MUSE and other beamforming algorithms are time-only models. In addition, we find that level-only models outperform time-only models in reverberant conditions, but do worse in non-reverberant enrionments. Intriguingly, DNNs trained with both time and level (purple) perform better than level-only models in reverberant environments, suggesting that DNNs are making use of both available cues, though likely relying more on level. Future studies will aim to better understand how DNNs and biological neural networks balance the relative use of these two cues in reverberant listening conditions.

## 2 Datasheet for VCL

### Motivation For Dataset Creation

**2.1 For what purpose was the dataset created? (e.g., Was there a specific task in mind? Was there a specific gap that needed to be filled?)**

Communication disorders affect more than 45 million individuals in the United States [5], however the neural mechanisms that underlay these disorders are poorly understood. The field of neuroscience has developed a number of animals models to study the neural basis of vocal communication, including social rodents, who naturally vocalize during social interactions. The VCL dataset and benchmark were established to address the problem of sound-source localization (SSL) and source attribution in the context of rodent social-vocal interactions. More colloquially, in a group of vocalizing individuals: **who said what**? Existing non-invasive approaches rely on classical signal processing algorithms, though their performance is contingent upon specialized acoustical hardware which is not ideal for next-generation neuroscience experiments (see main text Section 1 and 2.1). Therefore, an off-the-shelf SSL tool which works in standard reverberant laboratory environments is needed.

Advances in deep learning for machine audio are well poised to help address these challenges [2], however there is little interaction between the fields of machine audio and neuroscience. One of the functions of this dataset and benchmark release is to facilitate collaboration between these fields by formalizing the neuroscientific problem in terms of a machine learning problem. To this end, we acquired the first ever large-scale SSL dataset with ground truth labels for SSL and source attribution in rodent social interactions. In addition, we introduce two tasks which serve as benchmarks:

- **Task 1 - Sound Source Localization:** Compare the performance of classical sound source localization algorithms with deep neural networks.
- **Task 2 - Vocalization Attribution:** Assign vocalizations to individuals in a social dyad.

**2.2 What (other) tasks could the dataset be used for?**

Recent work[4] has shown that sound localization DNNs trained on synthetic data can be used to reliably predict sound sources in held-out real-world data. Indeed, preliminary experiments (see main text, Figure 2D) suggest that this is also true for the VCL dataset. Future experiments will explore how DNNs trained on synthetic only vs. real data augmented with synthetic data (with different proportions real:synthetic) affect performance.

This dataset can also be used to assess SSL generalization. For example, Supplementary Figure 1 shows that predictions from DNNs trained on a single stimulus class do not generalize well to other stimulus classes. Future tasks will explore different architectures and input embeddings that may facilitate generalization.

**2.3 Who created this dataset (e.g., which team, research group) and on behalf of which entity (e.g., company, institution, organization)?**

The dataset was created collaboratively between the Williams (NYU), Sanes (NYU), Schneider (NYU), Falkner (Princeton), and Murthy (Princeton) labs. Datasets from environments E1/E2 were acquired at NYU, E3 at Princeton.

**2.4 Who funded the creation dataset?**

This work was supported by the National Institutes of Health R34-DA059513 (AHW, DHS, DMS), National Institutes of Health R01-DC020279 (DHS), National Institutes of Health 1R01-DC018802 (DMS, REP), National Institutes of Health Training Program in Computational Neuroscience T90DA059110 (REP), New York Stem Cell Foundation (DMS), CV Starr Fellowship (BM), EMBO Postdoctoral Fellowship (BM), National Science Foundation Award 1922658 (CI).

**2.5 Any other comment?**

None.

### Dataset Composition

**2.6 What are the instances?(that is, examples; e.g., documents, images, people, countries) Are there multiple types of instances? (e.g., movies, users, ratings; people, interactions between them; nodes, edges)**

A single instance in the dataset consists of two features:

- Raw multi-channel audio data recorded from a microphone array.
- Ground truth source position (XY) of audio data.

**2.7 How many instances are there in total (of each type, if appropriate)?**

767,295 total sound events (instances) from 9 unique conditions. See main text, Table 1.

**2.8 What data does each instance consist of“Raw” data (e.g., unprocessed text or images)? Features/attributes? Is there a label/target associated with instances? If the instances related to people, are subpopulations identified (e.g., by age, gender, etc.) and what is their distribution?**

Each instance consists of raw multi-channel audio with a label indicating where in 2-dimensional space the sound came from. The sound itself comes from either speaker playback of vocalizations/sine sweeps or a real vocalizing animal.

**2.9 Is there a label or target associated with each instance? If so, please provide a description.**

Yes, the label for each instance is the 2-dimensional location of the sound source.

**2.10 Is any information missing from individual instances? If so, please provide a description, explaining why this information is missing (e.g., because it was unavailable). This does not include intentionally removed information, but might include, e.g., redacted text.**

None.

**2.11 Are relationships between individual instances made explicit (e.g., users’ movie ratings, social network links)? If so, please describe how these relationships are made explicit.**

Yes, samples were acquired from distinct environments, sources, and represent different stimulus types. See main text, Table 1 for additional detail.

**2.12 Does the dataset contain all possible instances or is it a sample (not necessarily random) of instances from a larger set? If the dataset is a sample, then what is the larger set? Is the sample representative of the larger set (e.g., geographic coverage)? If so, please describe how this representativeness was validated/verified. If it is not representative of the larger set, please describe why not (e.g., to cover a more diverse range of instances, because instances were withheld or unavailable).**

The dataset contains all possible instances.

**2.13 Are there recommended data splits (e.g., training, development/validation, testing)? If so, please provide a description of these splits, explaining the rationale behind them.**

We used a random 80:10:10 train, test, validation split with a predetermined random seed. The exact splits are available for download on the dataset website.

**2.14 Are there any errors, sources of noise, or redundancies in the dataset? If so, please provide a description.**

None that we are aware of.

**2.15 Is the dataset self-contained, or does it link to or otherwise rely on external resources (e.g., websites, tweets, other datasets)? If it links to or relies on external resources, a) are there guarantees that they will exist, and remain constant, over time; b) are there official archival versions of the complete dataset (i.e., including the external resources as they existed at the time the dataset was created); c) are there any restrictions (e.g., licenses, fees) associated with any of the external resources that might apply to a future user? Please provide descriptions of all external resources and any restrictions associated with them, as well as links or other access points, as appropriate.**

It is self-contained.

### Collection Process

**2.16 What mechanisms or procedures were used to collect the data (e.g., hardware apparatus or sensor, manual human curation, software program, software API)? How were these mechanisms or procedures validated?**

#### E1 Datasets

Audio, video, and speaker playback were synchronously acquired and triggered using custom python software, as per [7]. Four ultrasonic microphones (Avisoft CM16/CMPA48AAF-5V) connected to an Avisoft preamplifier were recorded by a National Instruments data acquisition device (PCI-6143) via BNC connection with a National Instruments terminal block (BNC-2110). The recording was controlled using the NI-DAQmx library (https://github.com/ni/nidaqmx-python) which wrote samples to disk at a 125 kHz sampling rate. Video (FLIR USB Backfly S) frames were externally triggered at 30 Hz via the National Instruments device and written to disk using the FLIR Spinnaker SDK, PySpin. Pre-computed wav files were stored on a Raspberry Pi 4B and played back using the “play” command from the SoX library. Audio signals from the Raspberry Pi were sent to a digital-to-analog converter (HiFiBerry DAC2 Pro), then to an amplifier (Tucker Davis Technologies SA-1), then finally to a speaker.

#### E2 Datasets

Audio and video were synchronously acquired in a similar fashion to E1 data, with two notable exceptions. First, instead of externally triggering camera frames, we recorded timestamps generated by a camera synchronization device at 30 Hz (e3 Vision Hub + Camera, White Matter LLC). Second, we recorded 8-channel audio using a different National Instruments configuration (PXI-6143 IO Module mounted in a PXIe-1071 chassis) at a sampling rate of 125 kHz. See **Hexapod Dataset** Section for information about audio playback in this environment.

#### E3 Datasets

Similarly to E1 and E2, audio and video were synchronously acquired using custom python scripts. 24 Avisoft microphones (CM16/CMPA) were connected to an Avisoft UltraSoundGate 1216H (analog-to-digital converter and pre-amplifier) and written to disk at a sampling rate of 250 kHz. Video (FLIR USB Backfly S) frames were externally triggered at 150 Hz using a Loopbio triggerbox. Audio and video were synchronized posthoc using 1.) TTLs sent every frame from the triggerbox to the Avisoft UltraSoundGate and 2.) by aligning simultaneous pulses sent from an Arduino to the Avisoft UltraSoundGate and to trigger LEDs detected by the camera. Audio playback was performed in the same manner as E1.

#### Speaker Dataset

Stimuli were played back with the system described in E1 using a Fountek NeoCD1 Ribbon Tweeter speaker. Upon playback of an audio signal, a TTL was sent from the Raspberry Pi to the National Instruments system and a timestamp for each playback was logged. The speaker was positioned facing towards the ceiling of the arena and moved in 2cm increments by hand to tile the arena floor uniformly. At each position in the arena, every stimulus class (see Supplementary Section 1.1) was played 10 times (180 stimuli total, per position). The ground truth position of the sound source was obtained using OpenCV.

#### Edison Dataset

A Bose SoundTrue Ultra in-ear earbud was magnetically mounted to the top of an Edison Robot v2 (https://meetedison.com/). The robot was programmed to perform a random walk using the EdPy library (https://www.edpyapp.com/) and using the speaker playback system detailed in E1, we played randomly sampled vocalizations from gerbil families [7] out of the earbud speaker every 50ms. The ground truth position of the sound source was obtained using supervised keypoint tracking with SLEAP [6].

#### Hexapod Dataset

The Hexapod dataset was generated using a Freenove Hexapod Robot Kit (https://freenove.com/). The kit features a 6 legged robot independently powered by two 18650 batteries and controlled by a custom microcontroller developed by Freenove. A HC-SR04 ultrasonic sensor is mounted at the front of the hexapod to ensure it avoids collisions with walls. Additionally, a second system is integrated into the hexapod, comprising a Raspberry Pi 4B, HifiBerry DAQ2 Pro, and a MAX9744 amplifier (Adafruit: 1752), all housed in 3D printed cases. This setup plays gerbil vocalizations through a Peerless by Tymphany tweeter speaker (DigiKey: OX20SC00-04-ND) mounted on a pan-tilt servo mechanism. This system is powered by a portable Anker power bank. The tweeter speaker is enclosed in a 3D printed box marked with multiple ArUco markers, enabling continuous tracking of the speaker’s 3D position. The robot is designed to operate untethered and is controlled wirelessly.

The control loop starts with the Raspberry Pi, which sends a signal to the hexapod microcontroller to start the robot’s gait. After a set number of steps, the hexapod microcontroller issues a stop signal to the Raspberry Pi and waits for another start signal before resuming movement. Meanwhile, the Raspberry Pi on receiving the stop signal transmits vocalizations through the HiFiBerry DAC and MAX9744 amp to the speaker. These vocalizations are emitted at various angles using the pan-tilt servo mechanism. The hexapod resumes movement only after completing a full cycle of speaker positions and vocalizations. A complete cycle involves four distinct pan servo positions—North, North East, East, South East, and South—and five tilt servo positions—0, 45, 90, 135, and 180 degrees. At each position, two sine frequencies, a sine sweep and 100 different vocalizations are played. The sine frequencies are used to detect the start of the audio sequence and the sine sweep is used to calibrate the room impulse response for simulations.

#### Earbud Datasets

Neodymium block magnets (K+J Magnetics: B222G-N52) were glued into a custom 3D printed housing, then affixed to the skulls of anesthetized gerbils or mice using dental cement or cyanoacrylate glue. A block magnet was then glued to an earbud (either Bose SoundTrue Ultra in-ear or Sony MDREX15LP in-ear) to allow for easy application and removal of the earbud to the animal’s head. After surgical recovery, animals were allowed to freely explore E1 (gerbils) or E3 (mice) with the head-mounted earbud. For gerbils, vocalizations were played back out of the earbud in the same fashion as the Edison Dataset. This process generated the GerbilEarbud-4M-E1 Dataset. For mice, a representative collection of ultrasonic vocalization types were used to generate a wav file containing 10,000 vocalizations that were played back over a period of 20 minutes (MouseEarbud-24M-E3 Dataset). The ground truth position of the sound source was obtained using supervised keypoint tracking with SLEAP.

#### Solo Gerbil Dataset

Adolescent gerbils respond robustly to playback of an 11-syllable, 2-second long sequence of ultrasonic vocalizations (REP, unpublished observations). We leveraged this behavior to generate ground-truth data by placing single adolescent gerbils in E1 and playing back the vocalization sequence to induce vocal responses from the isolated animal. We then tracked the XY position of the animal’s nose using SLEAP and extracted vocalization onset times using the supervised audio segmenter DAS [8].

#### Solo Mouse Dataset

Male mice vocalize in response to the smell of urine from sexually receptive females [3]. Prior to free exploration of E3 by a single male mouse, estrus females were first allowed explore the arena, thereby depositing their smell across the environment. Males reliably vocalized in isolation following this procedure. We then tracked the XY position of the animal’s nose using SLEAP and extracted vocalization onset times using the supervised audio segmenter DAS.

#### Gerbil/Mouse Dyad Datasets

Standard laboratory rodents naturally vocalize during social interactions, therefore gerbils or mice were recorded during dyadic social interactions. We tracked the XY position of the animals noses using SLEAP and extracted vocalization onset times using DAS. Although we do not have ground truth for which animal vocalized, we do know two possible locations of the sound source, therefore this dataset can be used for Task 2.

**2.17 How was the data associated with each instance acquired? Was the data directly observable (e.g., raw text, movie ratings), reported by subjects (e.g., survey responses), or indirectly inferred/derived from other data (e.g., part-of-speech tags, model-based guesses for age or language)? If data was reported by subjects or indirectly inferred/derived from other data, was the data validated/verified? If so, please describe how.**

The data were directly observable, as detailed above. In the case of datasets which used speaker playback, the onset times of vocalizations (i.e. instances) were automatically logged using a TTL sent to the data acquisition device. In the case of real rodent vocalizations where this was not possible, we used DAS, a deep-learning based audio segmenter, to extract the onset/offset times of vocalizations. To obtain the ground truth XY position of sound sources, we used SLEAP, a supervised keypoint tracking software, to infer the position of sound sources in the arena.

SLEAP and DAS are both supervised learning pipelines, therefore need sufficient and diverse training data to function appropriately. We validated the performance of these algorithms through iterative human-in-the-loop training, where the labeler trained a model with limited training data, then evaluated performance on heldout data. If the heldout performance was insufficient, the labeler added more training examples and retrained the model. This process was repeated until the models achieved desired performance.

**2.18 If the dataset is a sample from a larger set, what was the sampling strategy (e.g., deterministic, probabilistic with specific sampling probabilities)?**

N/A

**2.19 Who was involved in the data collection process (e.g., students, crowdworkers, contractors) and how were they compensated (e.g., how much were crowdworkers paid)?**

The dataset was acquired by graduate students and postdoctoral fellows at New York University and Princeton University. They are compensated by grants that support laboratories and/or individual grant funding (see Section 2.4).

**2.20 Over what timeframe was the data collected? Does this timeframe match the creation timeframe of the data associated with the instances (e.g., recent crawl of old news articles)? If not, please describe the timeframe in which the data associated with the instances was created.**

The datasets were acquired over a one year period from April 2023-May 2024.

**2.21 Data Preprocessing**

**2.22 Was any preprocessing/cleaning/labeling of the data done (e.g., discretization or bucketing, tokenization, part-of-speech tagging, SIFT feature extraction, removal of instances, processing of missing values)? If so, please provide a description. If not, you may skip the remainder of the questions in this section.**

Instances from experiment days were inspected to ensure that they contained XY positions that accurately labelled the sound source. In the event that a sound source position was mislabeled, we manually corrected the label(s) for those days, or used OpenCV to track the sound source instead of SLEAP (e.g. Speaker- 4M-E1 Dataset). Occasionally we omitted the entire instance if no sound source was visible in the camera view.

**2.23 Was the “raw” data saved in addition to the preprocessed/cleaned/labeled data (e.g., to support unanticipated future uses)? If so, please provide a link or other access point to the “raw” data.**

No, we only provided the curated dataset.

**2.24 Is the software used to preprocess/clean/label the instances available? If so, please provide a link or other access point.**

Yes, see https://github.com/neurostatslab/vocalocator.

**2.25 Does this dataset collection/processing procedure achieve the motivation for creating the dataset stated in the first section of this datasheet? If not, what are the limitations?**

Yes.

**2.26 Any other comments**

None.

### Dataset Distribution

**2.27 How will the dataset be distributed? (e.g., tarball on website, API, GitHub; does the data have a DOI and is it archived redundantly?)**

**Data is available at:** https://users.flatironinstitute.org/~atanelus **Code is available at:** https://github.com/neurostatslab/vocalocator The dataset DOI is: 10.5281/zenodo.11584391

**2.28 When will the dataset be released/first distributed? What license (if any) is it distributed under?**

As of this submission, the dataset is currently publicly available and distributed under a CC BY 4.0 license. The authors bear all responsibility in case of violation of rights.

**2.29 Are there any copyrights on the data?**

None.

**2.30 Are there any fees or access/export restrictions?**

None.

**2.31 Any other comments?**

None.

### Dataset Maintenance

**2.32 Who is supporting/hosting/maintaining the dataset?**

The Flatiron Institute Center for Computational Neuroscience and Scientific Computing Core.

**2.33 Will the dataset be updated? If so, how often and by whom?**

Yes, it will be updated as the neuroscience and ML community begin to use the datasets. Neuroscience labs will contribute their own datasets.

**2.34 How will updates be communicated? (e.g., mailing list, GitHub)**

GitHub + mailing list.

**2.35 If the dataset becomes obsolete how will this be communicated?**

Mailing list.

**2.36 Is there a repository to link to any/all papers/systems that use this dataset?**

N/A

**2.37 If others want to extend/augment/build on this dataset, is there a mechanism for them to do so? If so, is there a process for tracking/assessing the quality of those contributions. What is the process for communicating/distributing these contributions to users?**

A clear set of instructions for how to contribute a dataset will be included on the website. Contributors will be credited on the website/GitHub repo.

### Legal and Ethical Considerations

**2.38 Were any ethical review processes conducted (e.g., by an institutional review board)? If so, please provide a description of these review processes, including the outcomes, as well as a link or other access point to any supporting documentation.**

All procedures related to the maintenance and use of animals were approved by the Institutional Animal Care and Use Committee (IACUC) at New York University and Princeton University. All experiments were performed in accordance with the relevant guidelines and regulations.

**2.39 Does the dataset contain data that might be considered confidential (e.g., data that is protected by legal privilege or by doctorpatient confidentiality, data that includes the content of individuals non-public communications)? If so, please provide a description.**

No.

**2.40 Does the dataset contain data that, if viewed directly, might be offensive, insulting, threatening, or might otherwise cause anxiety? If so, please describe why**

No.

**2.41 Does the dataset relate to people? If not, you may skip the remaining questions in this section.**

N/A

**2.42 Does the dataset identify any subpopulations (e.g., by age, gender)? If so, please describe how these subpopulations are identified and provide a description of their respective distributions within the dataset.**

N/A

**2.43 Is it possible to identify individuals (i.e., one or more natural persons), either directly or indirectly (i.e., in combination with other data) from the dataset? If so, please describe how.**

N/A

**2.44 Does the dataset contain data that might be considered sensitive in any way (e.g., data that reveals racial or ethnic origins, sexual orientations, religious beliefs, political opinions or union memberships, or locations; financial or health data; biometric or genetic data; forms of government identification, such as social security numbers; criminal history)? If so, please provide a description.**

N/A

**2.45 Did you collect the data from the individuals in question directly, or obtain it via third parties or other sources (e.g., websites)?**

N/A

**2.46 Were the individuals in question notified about the data collection? If so, please describe (or show with screenshots or other information) how notice was provided, and provide a link or other access point to, or otherwise reproduce, the exact language of the notification itself.**

N/A

**2.47 Did the individuals in question consent to the collection and use of their data? If so, please describe (or show with screenshots or other information) how consent was requested and provided, and provide a link or other access point to, or otherwise reproduce, the exact language to which the individuals consented.**

N/A

**2.48 If consent was obtained, were the consenting individuals provided with a mechanism to revoke their consent in the future or for certain uses? If so, please provide a description, as well as a link or other access point to the mechanism (if appropriate).**

N/A

**2.49 Has an analysis of the potential impact of the dataset and its use on data subjects (e.g., a data protection impact analysis)been conducted? If so, please provide a description of this analysis, including the outcomes, as well as a link or other access point to any supporting documentation.**

N/A

**2.50 Any other comments?**

None.

## Footnotes

3 See, for example, Figure 1D in [54] for a distribution of inter-animal distances during natural social behavior.

